# Multiprotein Assemblies, Phosphorylation and Dephosphorylation in Neuronal Cytoskeleton

**DOI:** 10.1101/2023.06.21.545989

**Authors:** Natalya Kurochkina, Matthew R. Sapio, Michael J. Iadarola, Bradford E Hall, Ashok B. Kulkarni

## Abstract

Filament systems are comprised of fibrous and globular cytoskeletal proteins and are key elements regulating cell shape, rigidity, and dynamics. The cellular localization and assembly of neurofilaments depend on phosphorylation by kinases. The involvement of the BRCA1 (Breast cancer associated protein 1)/BARD1 (BRCA1-associated RING domain 1) pathways in Alzheimer disease (AD) is suggested by colocalization studies. In particular, BRCA1 accumulation within neurofibrillary tangles and colocalization with tau aggregates in the cytoplasm of AD patients implicates the involvement of mutant forms of BRCA1/BARD1 proteins in disease pathogenesis. The purpose of this study is to show that the location of mutations in the translated BARD1, specifically within ankyrin repeats, has strong correlation with the Cdk5 motifs for phosphorylation.

Mapping of the mutation sites on the protein’s three-dimensional structure and estimation of the backbone dihedral angles show transitions between the canonical helical and extended conformations of the tetrapeptide sequence of ankyrin repeats.

Clustering of mutations in BARD1 ankyrin repeats near the N-termini of the helices with T/SXXH motifs provides a basis for conformational transitions that might be necessary to ensure the compatibility of the substrate with active site geometry and accessibility of the substrate to the kinase.

Ankyrin repeats are interaction sites for phosphorylation-dependent dynamic assembly of proteins including those involved in transcription regulation and signaling, and present potential targets for the design of new drugs.

## Introduction

Filament systems are key elements regulating cell shape, rigidity, and molecular dynamics of receptor systems. The cytoskeleton is the site of assembly of fibrous and globular filamentous protein aggregates. Phosphorylation and dephosphorylation regulate cellular localization, assembly of complexes, and abnormal aggregation of neurofilaments (1, 2). Understanding microtubule dynamics, post-translational modifications, compartmental distribution, assembly of cellular components, and signaling linking neuronal dysfunction to neuropathies can lead to the development of new therapies.

The BRCA1 associated RING domain 1 (*BARD1*) gene has been studied extensively as a putative tumor suppressor and breast cancer susceptibility gene (3). To date the only known enzymatic function of BARD1 is as a ubiquitin ligase in complex with BRCA1. Heterodimerization of BARD1–BRCA1 via the RING domain is essential for the homologous recombination repair and transcriptional regulation functions of *BRCA1* (4). Together, this heterodimer is involved in DNA double-stranded break repair and cell-cycle regulation. Mutant forms of BRCA1/BARD1 have been strongly implicated in pathogenesis of breast cancer (5). Increasing evidence also supports the role of these proteins in other cancers, as well as pathological processes in other diseases. Involvement of the BRCA1/BARD1 heterodimer in Alzheimer’s disease (AD) is supported by several lines of evidence including the following: (1) BRCA1 accumulates within neurofibrillary tangles in the AD brains (6); (2) BRCA1 colocalizes with tau aggregates in the cytoplasm of AD patients (7).

In the present work, we examined common locations of missense mutations within the BARD1 ankyrin repeats. The mutations tend to localize near the N-termini of the helices, in which T/SXXH motifs were also identified. These T/SXXH motifs are potential substrates for kinases such as CDK5, based on predictions of similar kinase motifs from previous studies of this enzyme (8). Correlation between the location of the mutations and regularly positioned phosphorylation sites in comparison with the ankyrin repeat containing chemo-responsive ion channel TRPA1 provides a basis for conformational transitions that may be necessary for compatibility of the substrate with active site geometry, and ultimately accessibility of the substrate to the kinase.

## Materials and methods

Atomic coordinates of proteins determined by x-ray, cryoelectron microscopy and NMR were obtained from the Brookhaven Protein Databank (PDB). The following structural models given by their accession numbers were used: BARD1 (Accession: Q99728.2; PDB 1jm7 3c5r 2nte 2r1z 3fa2); TRPA1 (Accession: O75762; PDB 3j9p); CDK2 (Accession: P24941; PDB 1qmz); CDK5 (Accession: Q00535; PDB 1h4l), Estimation of backbone dihedral angles, interatomic distances, and molecular graphics were prepared using Jmol: an open-source Java viewer for chemical structures in 3D. http://www.jmol.org/.

## Results

BARD1 consists of several domains (Figure 1): RING (Really Interesting New Gene), Ankyrin repeats, and tandem BRCT (BRCA1 C-terminal repeat). The RING domain of BARD1 interacts with other proteins including BRCA1 (9). The BRCA1/BARD1 heterodimer localizes to the nucleus where it contributes to DNA repair and transcription activation. In the cytoplasm, BARD1 is implicated in apoptosis (10). The Ankyrin repeat-containing domain (ANK) interacts with proteins including the polyadenylation factor CstF-50 (11), p53, and BCL2 (12). At the C-terminus of the BARD1 there are two BRCT domains that interact with other proteins in phosphorylation dependent manner (13). Crystal and NMR structures are determined for these domains: RING (PDB 1jm7), ANK (PDB 3c5r), and BRCT (PDB 2nte 2r1z 3fa2).

**Figure 1.**
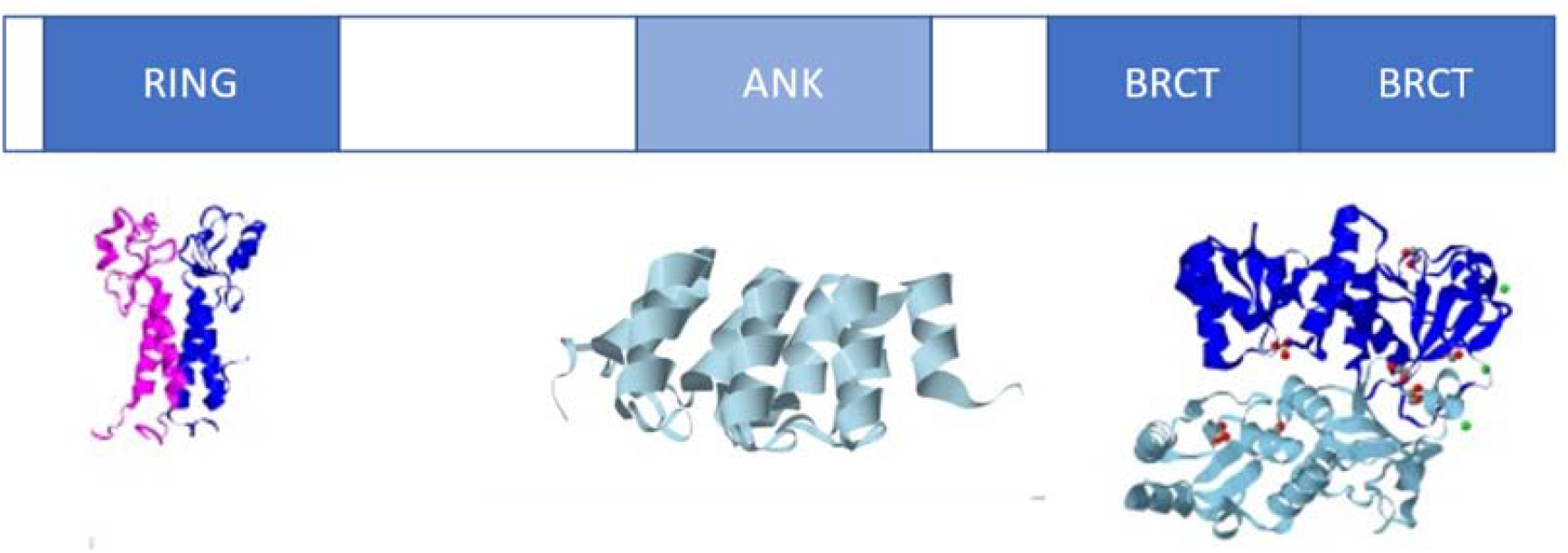
BARD1: Domain structure

Each unit of an individual ankyrin repeat consists of a pair of α-helices and a β-hairpin (Figure 2). Each inner row helix A forms an antiparallel interface with the outer row helix B and two parallel interfaces, one with preceding and one with the following helix in the row. As a result, AB, AA’, and AA’’ interfaces are observed in the inner row and, correspondingly, BB’ and BB’’ in the outer row. From the assignment of positions, a, d, e, and g, which are interface positions on the helix, we can see that inversion of the contacts at the helix-helix interfaces of the inner versus outer row results in the change of the direction of the row. This makes the two rows compatible when they come together in an antiparallel manner (14, 15). When amino acid sequences of the repeats are superimposed, substantial conservation at the amino acid level is consistent with a known consensus sequence (Figure 2) (16).

**Figure 2.**
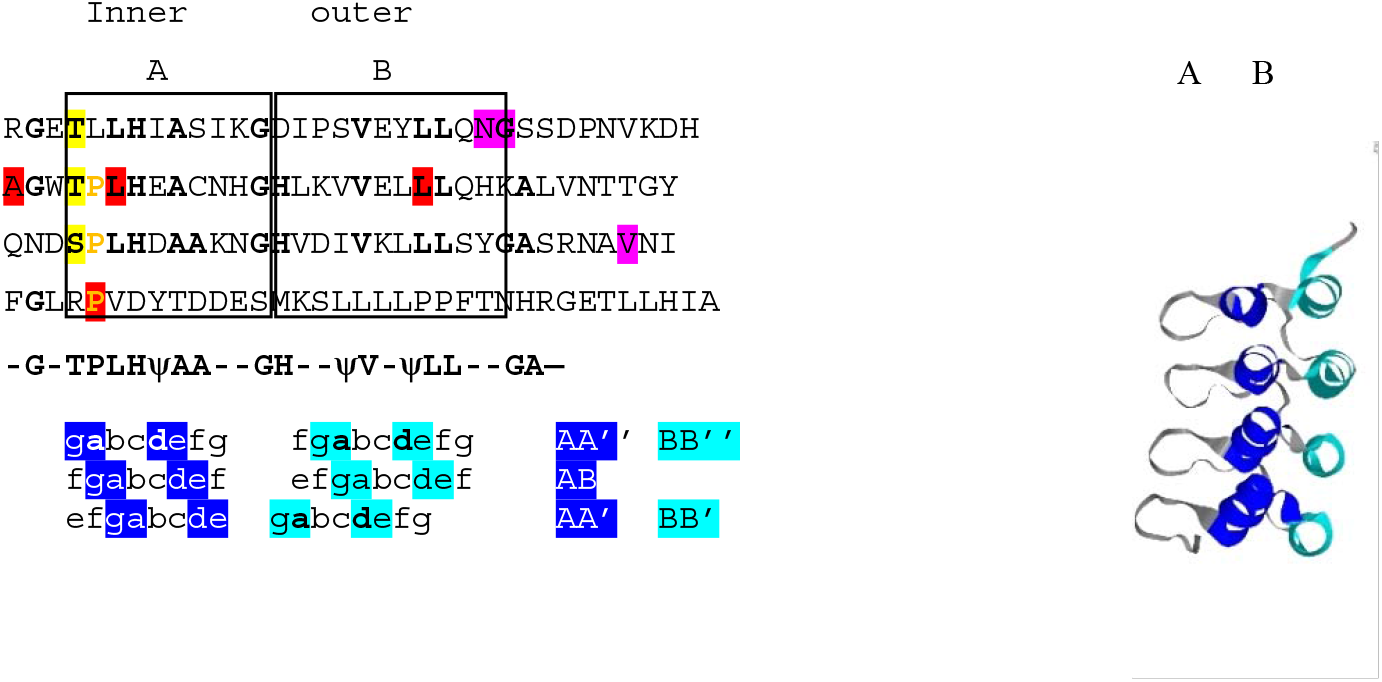
BARD1 mutations (functional X and nonfunctional X as in [5] and phospho-sites (T) in ankyrin repeats. Helices A and B boxed. Say what the BRCT domain is here.

More than 76 *BARD1* missense mutations were found in cancer patients. Out of these 76 mutants, six have a significantly higher loss of heterozygosity (LOH) and, therefore, increased likelihood of being pathogenic. Three of these mutations are located in the ankyrin repeats domain: 2 mutations in repeat 1 and 1 mutation in repeat 3, mainly at the C-terminus of helix B (V523A, N450H, G451fs). These mutations do not result in loss of function (5) (Figure 3). Another group of mutations contains those that are found in variants deficient in homologous DNA repair (HDR): A460T, L480S, and P530L. These mutations are located mainly at the N-terminus of helix A, and only one mutation at the C-terminus of helix B.

**Figure 3.**
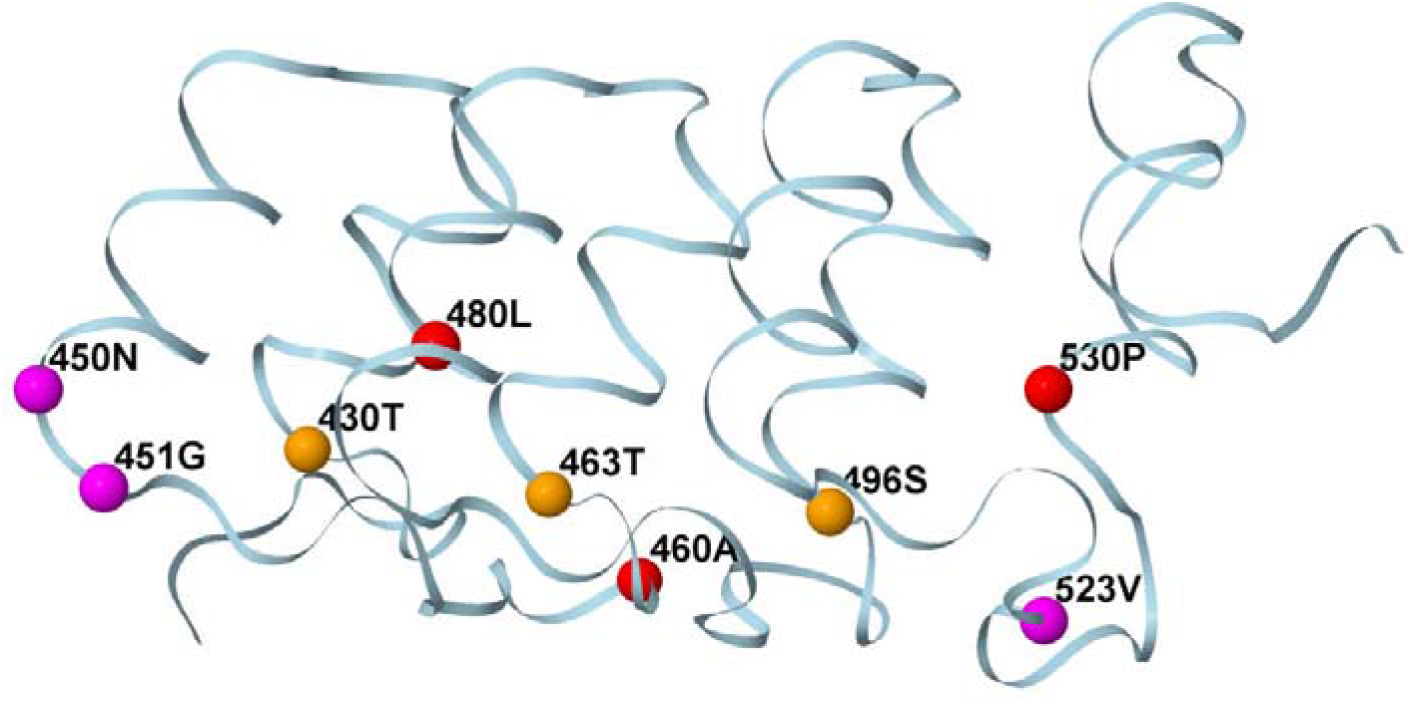
Structure of BARD1 ankyrin repeats domain which contains three repeats. Putative CDK5 phosphorylation sites (yellow) in helix 1 of each repeat appear to be in the proximity of pathogenic mutations (magenta and red residues) which are at the C-terminus of the helix 2 of repeats 1 and 3.

Similar to TRPA1 repeats, BARD1 repeats contain Ser/Thr followed by a Pro/Ala at the sites of TRPA1 phosphorylation by CDK5. Mutated sites in the BARD1 protein show strong correlation with the location of potential CDK5 phosphorylation sites. Even those mutations that are distant from phosphosites in amino acid sequence, are proximal in three-dimensional structure since N-terminus of helix A is near C-terminus of helix B. In three dimensions, both groups of mutations cluster so that amino acid substitutions in the BARD1 protein and potential phospho-sites are proximal.

In TRPA1, we know that Ser/Thr at the N-terminal positions of the repeats are indeed phosphorylated by CDK5 (8). TRPA1, transient receptor potential ankyrin 1, is a member of TRP family. The TRP family of channels is implicated in sensing a variety of stimuli including temperature, chemicals, pH, and the concentrations of various endogenous lipids (17). The TRPA1 molecule contains 17 ankyrin repeats, 6 of which contain a consensus sequence consistent with the substrate specificity of CDK5. The 6 consensus sites are repeats number 2, 3, 6, 11, 12, and 13 (using the numbering in Cdk5 from mouse) (8). All of them contain the Ser/ThrXXHis tetrapeptide motif that is common to approximately 41% of ankyrin repeats(18). The structure of TRPA1 has been previously determined by cryoelectron microscopy (PDB 3j9p), as well as theoretical modelling of the N-terminal ankyrin repeats (8). It is evident from the structure that CDK5 phosphorylation sites coincide with the N-terminal positions carrying tetrapeptide motif sequences (Figure 4). Conformation of the N-terminus of helix A of each repeat is characterized by ϕ,ψ dihedral angles of the backbone (Table). The site of the tetrapeptide motif of helix A is subject to conformational transitions. One set of conformations is consistent with canonical α-helix as in R12 of TRPA1. Another set of conformations is consistent with an extended conformation as in BARD1 and R13 of TRPA1. The helix and extended conformations are common for this tetrapeptide sequence. The extended conformation is consistent with conformation of the peptide substrate bound to the active site of the CDK enzyme. Therefore, we conclude that the N-terminus of helix A, the site of the tetrapeptide motif, is the site of conformational changes.

**Table.** Comparison of the conformations at the N termini of the ankyrin repeats in BARD1, TRPA1, and Cdk2.

| **BARD1 R1** |  |  | **BARD1 R2** |  |  | **TRPA1 R12** |  |  | **TRPA1 R13** |  |  | **Cdk2 substrate** |  |  |
| --- | --- | --- | --- | --- | --- | --- | --- | --- | --- | --- | --- | --- | --- | --- |
| **PDB 3c5r** |  |  | **PDB 3c5r** |  |  | **PDB 3j9p** |  |  | **PDB 3j9p** |  |  | **PDB 1qmz** |  |  |
| **aa** | **φ** | **ψ.** | **aa** | **φ** | **ψ.** | **aa** | **φ** | **ψ.** | **aa** | **φ** | **ψ.** | **aa** | **φ** | **ψ.** |
|  |  |  |  |  |  | K447 | 114 | -110 | M483 | 160 | -158 | A4 | 161 | -85 |
| **T430** | **169** | **-85** | **T463** | **158** | **-99** | **S448** | **-44** | **-102** | **T484** | **152** | **-123** | **S5** | **137** | **-66** |
| L431 | -38 | -54 | P464 | -36 | -50 | P449 | -37 | -52 | P485 | -37 | -48 | P6 | 135 | -63 |
| L432 | -45 | -62 | L465 | -44 | -64 | L450 | -42 | -50 | L486 | -41 | -58 | R7 | 121 | -94 |
| H433 | -52 | -54 | H466 | -46 | -56 | H451 | -50 | -55 | H487 | -58 | -69 |  |  |  |

**Figure 4.**
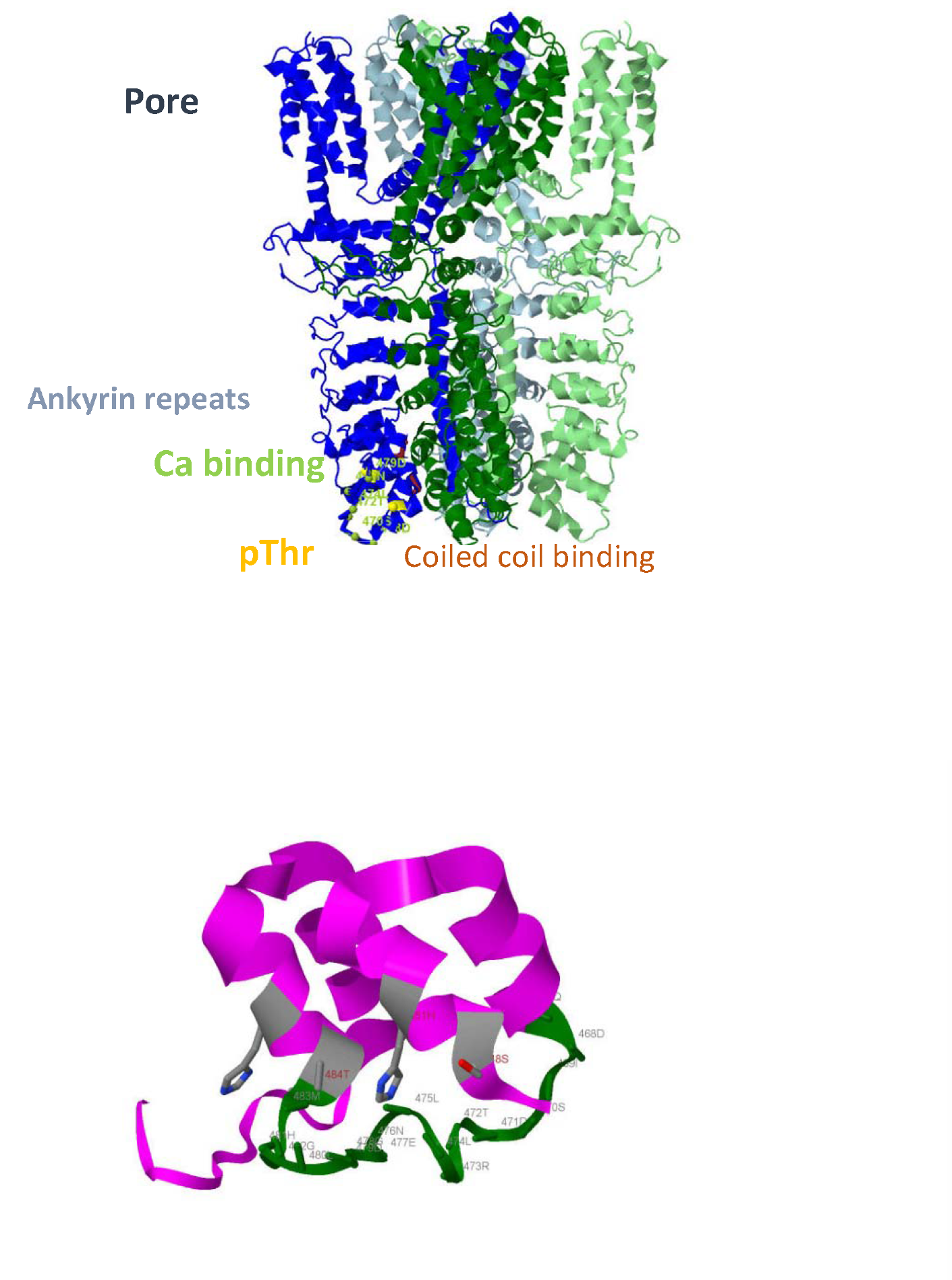
TRPA1 phosphorylation at structurally equivalent positions of ankyrin repeats. TRPA1 structure (top). Two ankyin repeats, Ank12 and Ank13 (bottom). Calcium binding site (green). At high Ca2+ concentrations the site in its calcium bound form screens the Thr/Ser from the solvent and kinase; at low Ca2+ concentrations Thr/Ser is more accessible.

CDK5 is a proline-directed serine/threonine protein kinase (19). It functions in brain development and neuronal differentiation. Expression of CDK5 is not restricted to neurons. Active CDK5 complexes primarily reside on the cell membrane (targeted by myristoylated p35 and p39). Dysregulation of CDK5 is associated with Alzheimer’s disease pathogenesis (20). Dysfunction of CDK5 links to cell cycle reentry and elevated DNA double strand breaks. Like other CDKs, CDK5 regulates cell cycle progression and shares substrate specificity. Unlike other CDKs, CDK5 is active in adult brain and requires p35 for its activity (noncyclin).

The T/SXXH motif at the N terminus of helix A is characteristic of ankyrin repeats; this motif stabilizes the helix. Within this motif, the S/T hydroxyl group forms a hydrogen bond between the amide nitrogen of the backbone and the Nδ1 atom of the His side chain three residues apart. In TRPA1 repeats, His/Thr bifurcated hydrogen bonds are formed between the Nδ1 atom of His and both the backbone N−H and sidechain Oγ-H of threonine within the conserved T/SXXH motif of ankyrin repeats.

Changes in hydrogen bonding distances show that the N-terminus of the helix undergoes conformational transitions. In the repeat in the BARD1 protein, the hydrogen bond distance is to 2.9 Å, a characteristic distance between donor and acceptor. In TRPA1 repeats, the distance between hydrogen bond donor and acceptor is longer (5.2 to 6.1 Å; Figure 4). This provides evidence that the N terminus of the helix exhibits significant rearrangements. Furthermore, these rearrangements may be necessary to assure compatibility of the peptide with active site geometry. Another important feature of TRPA1 repeats is that T/SXXH sequence is located in the interior of the protein and is protected from solvent or kinase. Beta-hairpins or insertions such as a Ca^2+^ binding site, EF hand, in R13 create a barrier. Calcium binding makes the TXXH motif inaccessible, whereas in the calcium-free conformation, it becomes accessible.

If we compare conserved features of TRPA1 and BARD1 repeats, we can see that in TRPA1, at high Ca2+ concentrations the calcium bound EF hand protects the Thr/Ser sites from the solvent or kinase; at low Ca2+ concentrations, the EF hand releases calcium and the Thr/Ser residues become more accessible. In BARD1, the mutated amino acids in the proximity of Thr/Ser sites increase or decrease accessibility of Thr/Ser within the T/SXXH motif. Both molecules, BARD1 and TRPA1, exhibit canonical helical or extended conformations and rearrangements can occur around the TXXH site. The motif is protected by hairpins and insertions. Proximal β-hairpins, calcium-binding sites and mutations modulate accessibility of the substrate to kinase.

## ASSOCIATED CONTENT

BARD1: Q99728.2

TRPA1: O75762

CDK2: P24941

CDK5:: Q00535

## Discussion

Ankyrin repeats provide a scaffold for the assembly of various proteins (21). Sites of clustering of mutations in BARD1 ankyrin repeats near the N-termini of the helices with T/SXXH motifs bring our attention to the mechanisms that are affected by structure modifications. These modifications are of potential physiological importance in both cancer and neurodegenerative pathways. Particularly, we examined BARD1 in comparison with TRPA1, a protein with several phosphorylated ankyrin repeat domains. This comparison highlights the correlation between the location of the mutations and regularly positioned phosphorylation sites which suggests that conformational transitions and hydrogen bonding patterns might be necessary to assure compatibility of the substrate with active site geometry and accessibility of the substrate to the kinase. This contributes to existing knowledge of the molecular interaction between the kinase active site and ankyrin repeat domains and suggests additional molecular biological investigations should be performed to clarify this correlation.

Studies of dynamic structural changes, posttranslational modifications, and their influence on the function of the molecules may provide a foundational understanding for investigating neuronal dysfunction and neuropathies. This ultimately has implications for the development of new therapies directed towards this molecular target.

In summary, BARD1 is an important protein in the pathogenesis of Alzheimer’s Disease, and further protein chemistry studies will be important to clarify molecular mechanisms predicted from its structure. Ankyrin repeats are interaction sites of phosphorylation-dependent dynamic assembly of proteins and nucleic acids including those involved in transcription regulation and signaling, and present promising targets for the design of new drugs.

## Conflicts of interest

The authors declare that they have no conflicts of interest.

## Author Contributions

NK: Conceptualization, Investigation, Writing-original draft, Writing-review & editing. MS: Conceptualization, Investigation, Writing-original draft, Writing-review & editing. MI: Conceptualization, Investigation, Writing-original draft, Writing-review, Validation & editing., BH: Conceptualization, Investigation, Writing-original draft, Writing-review & editing., AK: Validation, Writing-review & editing,

## Funding Sources

Study was funded by the research funds of The School of Theoretical and Modeling and partially supported by the intramural research program of the NIH Clinical Center and National Institute of Dental and Craniofacial Research.

## ABBREVIATIONS

CCR2: CC chemokine receptor 2
CCL2: CC chemokine ligand 2
CCR5: CC chemokine receptor 5
TLC: thin layer chromatography.
BARD1: BRCA1 associated RING domain 1
BRCA1: Breast cancer associated protein 1
BRCT: BRCA1 C-terminal
CDK5: Cyclin-dependent kinase 5
RING: Really Interesting New Gene
TRPA1: Transient receptor potential ankyrin 1 ion channel.

## Notes

### Competing Interest Statement

The authors have declared no competing interest.

## REFERENCES

1. Boumil, E. F., Vohnoutka, R., Lee, S., Pant, H., and Shea, T. B. (2018). Assembly and turnover of neurofilaments in growing axonal neurites. Biol. Open 7: bio028795. doi: 10.1242/bio.028795

2. Kurochkina N, Bhaskar M, Yadav SP, Pant HC. Front Mol Neurosci. 11, 373 (2018).

3. Tarsounas M, Sung P. The antitumorigenic roles of BRCA1-BARD1 in DNA repair and replication. Nat Rev Mol Cell Biol. 2020 May;21(5):284–299. doi: 10.1038/s41580-020-0218-z. Epub 2020 Feb 24. PMID: 32094664; PMCID: PMC7204409.1.

4, Irminger-Finger I, Jefford CE. Is there more to BARD1 than BRCA1? Nat Rev Cancer. 2006 May;6(5):382–91. doi: 10.1038/nrc1878. PMID: 16633366.

5. Adamovich AI, Banerjee T, Wingo M, Duncan K, Ning J, Martins Rodrigues F, Huang KL, Lee C, Chen F, Ding L, Parvin JD. Functional analysis of BARD1 missense variants in homology-directed repair and damage sensitivity. PLoS Genet. 2019 Mar 29;15(3): e1008049. doi: 10.1371/journal.pgen.1008049. PMID: 30925164; PMCID: PMC6457558.

6. Nakamura M, Kaneko S, Dickson DW, Kusaka H. Aberrant Accumulation of BRCA1 in Alzheimer Disease and Other Tauopathies. J Neuropathol Exp Neurol. 2020 Jan 1;79(1):22–33. doi: 10.1093/jnen/nlz107. PMID: 31750914.

7. Kurihara M, Mano T, Saito Y, Murayama S, Toda T, Iwata A. Colocalization of BRCA1 with Tau Aggregates in Human Tauopathies. Brain Sci. 2019 Dec 20;10(1):7. doi: 10.3390/brainsci10010007. PMID: 31861888; PMCID: PMC7016802.

8. Hall BE, Prochazkova M, Sapio MR, Minetos P, Kurochkina N, et al. Sci Rep. 8, 1177 (2018).

9. Brzovic PS, Rajagopal P, Hoyt DW, King MC, Klevit RE. Structure of a BRCA1-BARD1 heterodimeric RING-RING complex. Nat Struct Biol. 2001 Oct;8(10):833–7. doi: 10.1038/nsb1001-833. PMID: 11573085.

10. Hawsawi YM, Shams A, Theyab A, Abdali WA, Hussien NA, Alatwi HE, Alzahrani OR, Oyouni AAA, Babalghith AO, Alreshidi M. BARD1 mystery: tumor suppressors are cancer susceptibility genes. BMC Cancer. 2022 Jun 1;22(1):599. doi: 10.1186/s12885-022-09567-4. PMID: 35650591; PMCID: PMC9161512.

11. Edwards RA, Lee MS, Tsutakawa SE, Williams RS, Nazeer I, Kleiman FE, Tainer JA, Glover JN. The BARD1 C-terminal domain structure and interactions with polyadenylation factor CstF-50. Biochemistry. 2008 Nov 4;47(44):11446–56. doi: 10.1021/bi801115g. Epub 2008 Oct 9. Erratum in: Biochemistry. 2009 May 12;48(18):4008. Nazeer, I [added]; Kleiman, F E [added]. PMID: 18842000; PMCID: PMC2654182.

12. Tembe V, Henderson BR. BARD1 translocation to mitochondria correlates with Bax oligomerization, loss of mitochondrial membrane potential, and apoptosis. J Biol Chem. 2007 Jul 13;282(28):20513–22. doi: 10.1074/jbc.M702627200. Epub 2007 May 17. PMID: 17510055.

13. Birrane G, Varma AK, Soni A, Ladias JA. Crystal structure of the BARD1 BRCT domains. Biochemistry. 2007 Jul 3;46(26):7706–12. doi: 10.1021/bi700323t. Epub 2007 Jun 6. PMID: 17550235.

14. Kurochkina NA, Iadarola MJ. Helical assemblies: structure determinants. J Theor Biol. 2015 Mar 21; 369:80–84. doi: 10.1016/j.jtbi.2015.01.012. Epub 2015 Jan 19. PMID: 25613414.

15. N. Kurochkina. Protein Structure and Modeling. Springer 2019.

16. Mosavi LK, Cammett TJ, Desrosiers DC, Peng ZY. The ankyrin repeat as molecular architecture for protein recognition. Protein Sci. 2004 Jun;13(6):1435–48. doi: 10.1110/ps.03554604. PMID: 15152081; PMCID: PMC2279977.

17. López-Requena A, Boonen B, Van Gerven L, Hellings PW, Alpizar YA, Talavera K. Roles of Neuronal TRP Channels in Neuroimmune Interactions. In: Emir TLR, editor. Neurobiology of TRP Channels. Boca Raton (FL): CRC Press/Taylor & Francis; 2017. Chapter 15. PMID: 29356476.

18. Preimesberger MR, Majumdar A, Aksel T, Sforza K, Lectka T, Barrick D, Lecomte JT. Direct NMR detection of bifurcated hydrogen bonding in the α-helix N-caps of ankyrin repeat proteins. J Am Chem Soc. 2015 Jan 28;137(3):1008–11. doi: 10.1021/ja510784g. Epub 2015 Jan 17. PMID: 25578373; PMCID: PMC4311973.

19. ,Shupp A, Casimiro MC, Pestell RG. Biological functions of CDK5 and potential CDK5 targeted clinical treatments. Oncotarget. 2017 Mar 7;8(10):17373–17382. doi: 10.18632/oncotarget.14538. PMID: 28077789; PMCID: PMC5370047.k

20. Batra S, Jahan S, Ashraf A, Alharby B, Jawaid T, Islam A, Hassan I. A review on cyclin-dependent kinase 5: An emerging drug target for neurodegenerative diseases. Int J Biol Macromol. 2023 Mar 1; 230:123259. doi: 10.1016/j.ijbiomac.2023.123259. Epub 2023 Jan 12. PMID: 36641018.

21. Stevens SR, Rasband MN. Pleiotropic Ankyrins: Scaffolds for Ion Channels and Transporters. Channels (Austin). 2022 Dec;16(1):216–229. doi: 10.1080/19336950.2022.2120467. PMID: 36082411; PMCID: PM

